# Body weight modifies the efficacy of lifespan-extending interventions in *ad libitum*-fed mice

**DOI:** 10.64898/2026.11.10.765234

**Authors:** Paul-Edouard Bouvet, Linda Partridge

## Abstract

Geroprotective interventions are often assessed by extension of lifespan in *ad libitum*-fed mice. Increased lifespan could therefore reflect rescue of adverse phenotypes of unhealthy controls. We analysed body weight and survival for 33,780 mice in the Interventions Testing Program (ITP) to test whether treatment effects varied with body weight. In males, each added standard deviation of attained adult weight made treatment 6% more favourable (HR 0.94), more apparent for interventions that lowered body weight (HR 0.90). Median lifespan increase rose from 6.2% in mice in the lightest quartile to 20.2% in the heaviest pre-dosing weight quartile among those experiments, mainly because control survival declined more steeply with weight than that of the intervention group. The effect of body weight was consistent across 13 of 14 compounds and reflected weight attained during adulthood rather than in early life. Weight-dependence of intervention effects was weak in ITP females, possibly because median survival of controls exceeded that of every sampled non-ITP female control group. Amelioration of weight-associated risk is therefore implicated as a major contributor to lifespan increases relative to heavy, short-lived controls. The geroprotective effects of these interventions may thus be less apparent in metabolically healthier mice or humans.

## Main

Mouse lifespan studies are generally conducted under *ad libitum* feeding and they express efficacy as a difference from a control group. The biological meaning of that contrast therefore depends in part on the condition of the control group. An intervention could increase lifespan by raising the survival ceiling for healthy mice or by reversing unhealthy features of the controls. Continuous access to food under sedentary housing can induce adiposity, insulin resistance and shortened survival relative to a metabolically healthy state.^1^ Reversal of these adverse phenotypes could therefore generate an increase in survival.

This distinction matters for translation. Geroprotectors could potentially be applied to the general human adult population, but their efficacy could vary between individuals according to their initial health status. Protective effects driven largely by reversal of weight-associated pathology in heavy mice may translate less readily to healthy-weight mice and humans. Experimental evidence supports this kind of dependence. In male C57BL/6 mice, resveratrol increased remaining lifespan by about 25% under a high-calorie diet, restoring survival to near that of controls on standard diet, but did not extend lifespan on that diet. The SIRT1 activator SRT1720 increased mean lifespan by 21.7% under a high-fat diet against 8.8% on a standard diet.^2,3^ The nutritional status of the mice thus had a major influence on the extension of life by these compounds.

Across interventions, greater weight reduction has been associated with larger lifespan increases.^4^ Smaller treatment effects have been reported with longer-lived control cohorts in the National Institute on Aging (NIA) Interventions Testing Program (ITP)^5^ and an inverse association between young-adult body mass and lifespan has also been reported within UM-HET3 mice, particularly in males.^6,7^ However, these observations do not establish whether the effectiveness of an intervention differs between lighter and heavier animals, which requires an individual-level treatment-by-weight analysis, an approach established in clinical trials for assessing treatment-effect heterogeneity.^8^

The ITP provides a comprehensive resource for this type of analysis. UM-HET3 mice are genetically heterogeneous, individuals are allocated randomly to intervention and control groups, and cohorts are replicated across three sites using standardized husbandry and survival ascertainment.^9^ GeneNetwork, a public phenotype database, adds harmonized longitudinal, maternal and genetic traits.^10^ Together, these resources distinguish early-life weight from attained adult weight and allow us to test within each experimental cohort whether treatment effects vary with either.

In this study, we sought to determine whether treatment effects on lifespan increase across the pre-treatment weight distribution, whether the pattern reflects early-life or attained adult weight, and whether sex, genetic architecture or husbandry help to explain the heterogeneity.

## Results

### Treatment efficacy rises with adult weight in male mice

We first examined whether individual body weight affected the magnitude of the survival response to interventions (Fig. 1a). Mice randomly assigned to each intervention regimen were compared with concurrent controls from the same cohort and site. Each such comparison, or experiment, is one intervention at a given dose and starting age, tested in one cohort, where a cohort is a single yearly breeding group of UM-HET3 mice distributed across the three testing sites. The same compound could contribute several experiments through different cohorts or dosing regimens. This gave 92 experiments with males (17,943 mice) and 88 experiments with females (15,837 mice) from 17 cohorts. Because mice continue to gain weight through much of adulthood, and later weights increasingly reflect treatment and weight loss before death, our analyses used the first available adult weight measurement.

**Figure 1.**
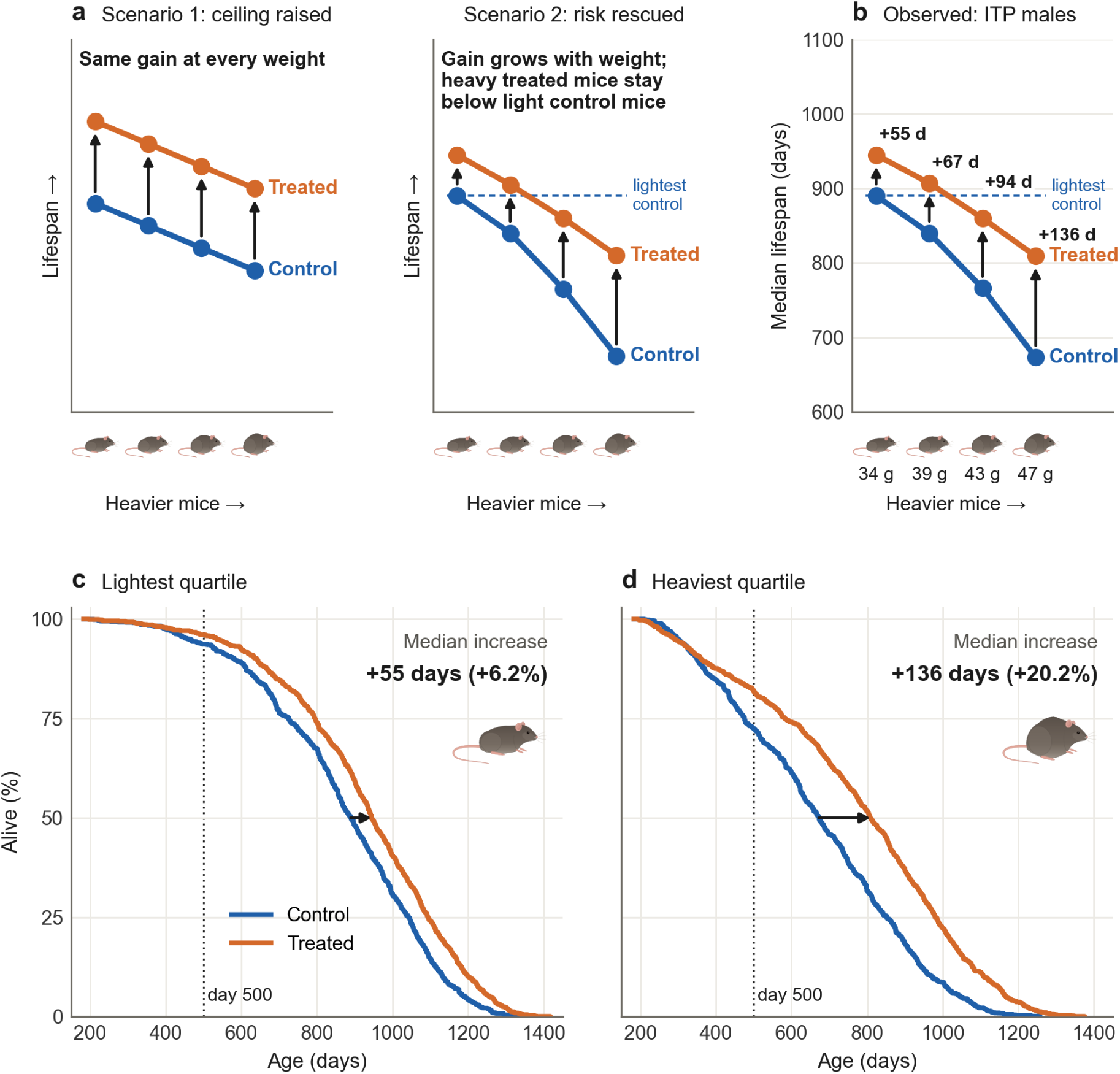
Lifespan increase is larger in heavier mice. a, Two ways in which the lifespan increase could depend on weight (schematic). In scenario 1, treatment raises lifespan by the same amount at every weight; in scenario 2, it mainly offsets the excess mortality of heavier controls, so the increase grows with weight and heavy treated mice remain shorter-lived than light controls (dashed line). b–d, Experiment- and site-standardized Kaplan–Meier survival across 30 weight-lowering experiments in males (8,137 unique mice), conditional on survival to six months. b, Median lifespan of control and treated mice in each pre-treatment weight quartile, with mean control weight below and the median increase in days next to each arrow; the dashed line marks the lightest-quartile control median. c,d, Control and treated mice in the lightest and heaviest pre-treatment weight quartiles, respectively. Black arrows show median lifespan increases; the dotted line marks day 500. The same within-experiment quartile definitions and standardization are used in Fig. 2b and Table 2.

In males, lifespan extension increased with weight relative to others in the same experiment. Each additional standard deviation of weight made the treatment-versus-control hazard ratio approximately 6% more favourable (HR 0.94, 95% CI 0.91–0.98; P = 0.0024; Fig. 2a; Table 1).

**Figure 2.**
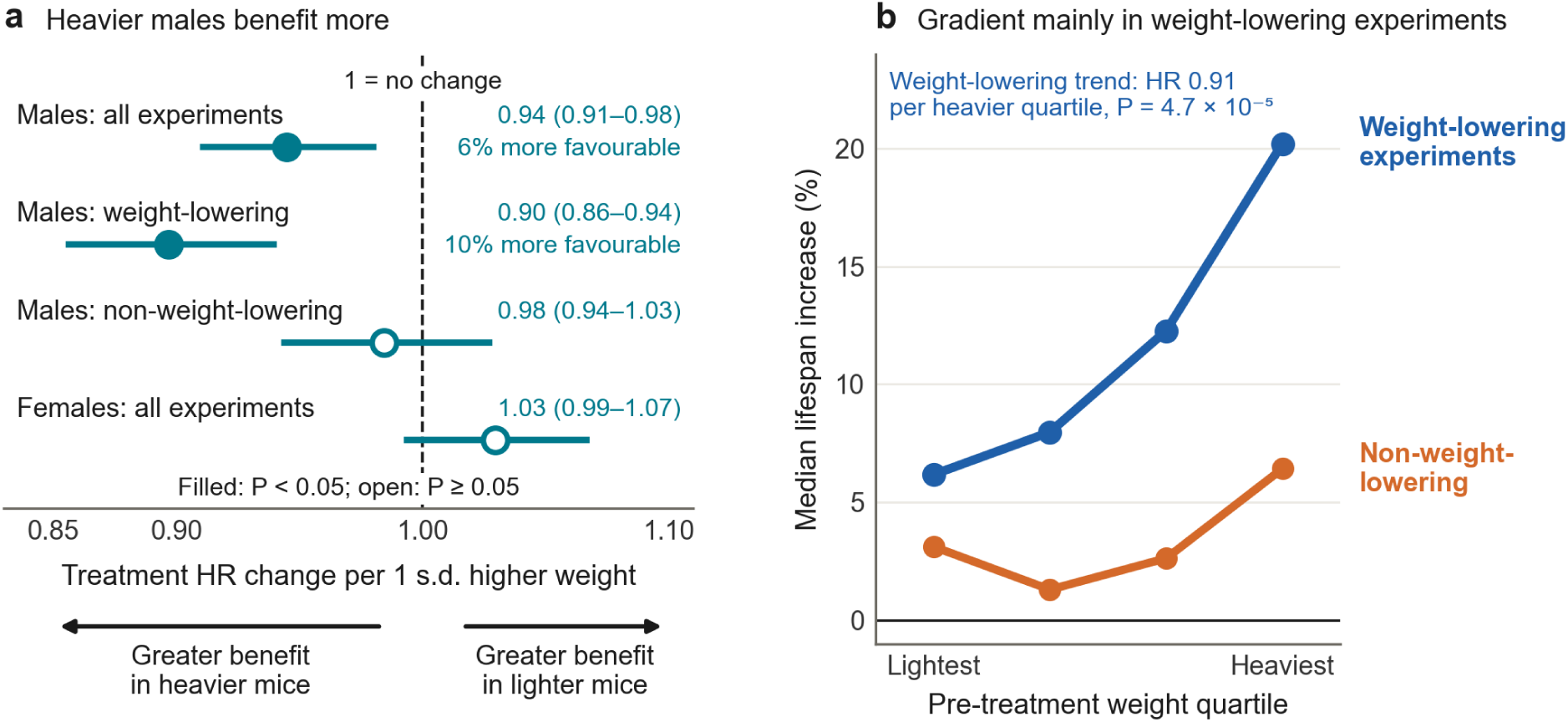
Weight dependence of the lifespan increase is concentrated in males and in weight-lowering interventions. a, Treatment-by-weight interaction HRs per 1 s.d. higher weight, with 95% CIs (Table 1); values below 1 indicate greater benefit in heavier mice. b, Median lifespan increase from the lightest to the heaviest pre-dosing weight quartile in 30 weight-lowering experiments in males (8,137 unique mice; experiment- and site-standardized Kaplan–Meier medians) and in 30 non-weight-lowering experiments in males (site-pooled medians). Quartiles were formed within each randomized group of each site-level comparison; the ordinal treatment-by-quartile trend in weight-lowering experiments was HR 0.91 (95% CI 0.87–0.95; P = 4.7 × 10⁻⁵; left-truncated Cox model with mouse-clustered variance, two-sided Wald test).

**Table 1.**
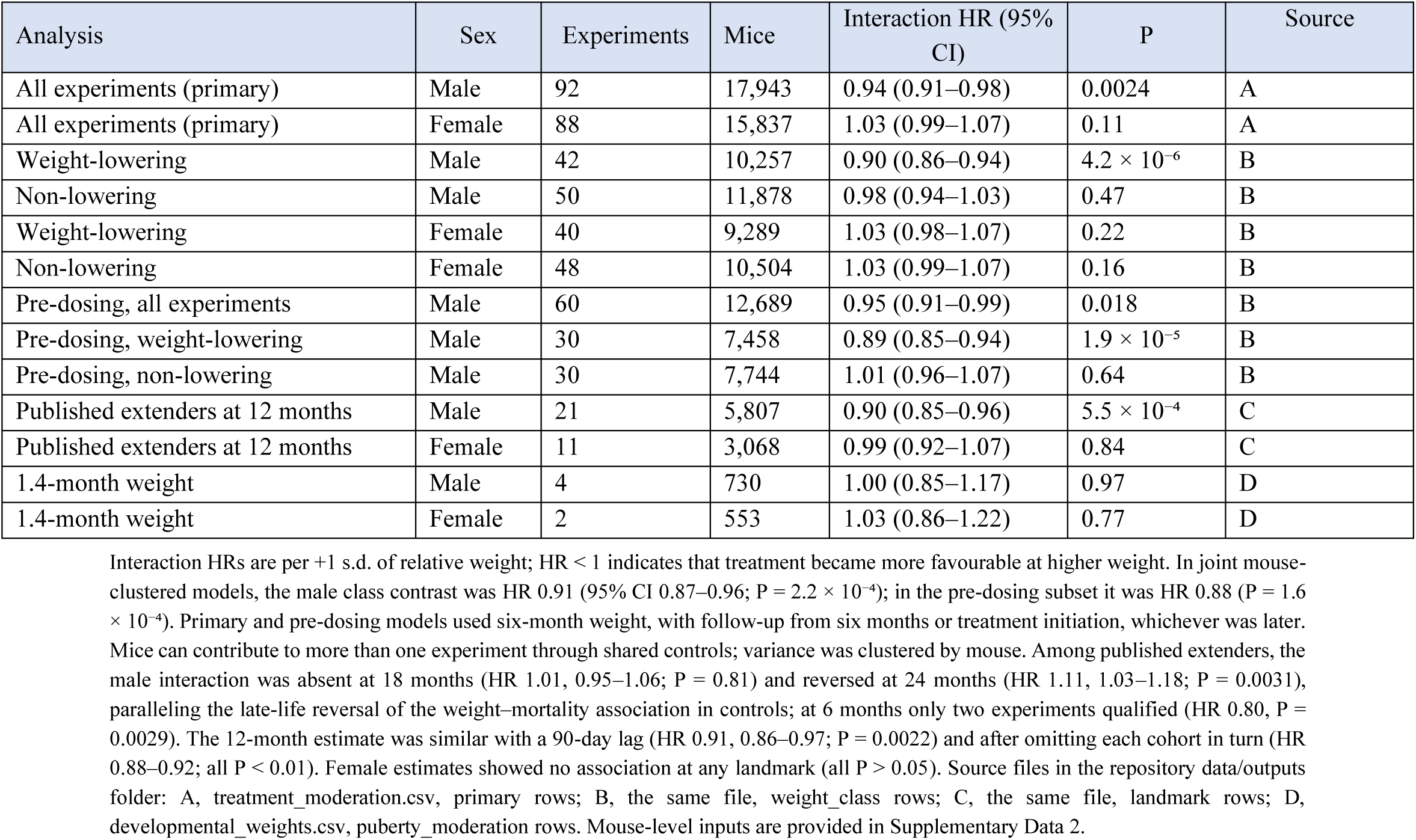
Treatment-by-weight moderation estimates.

| Analysis | Sex | Experiments | Mice | Interaction HR (95% CI) | P | Source |
|---|---|---|---|---|---|---|
| All experiments (primary) | Male | 92 | 17,943 | 0.94 (0.91–0.98) | 0.0024 | A |
| All experiments (primary) | Female | 88 | 15,837 | 1.03 (0.99–1.07) | 0.11 | A |
| Weight-lowering | Male | 42 | 10,257 | 0.90 (0.86–0.94) | 4.2 × 10<sup>−6</sup> | B |
| Non-lowering | Male | 50 | 11,878 | 0.98 (0.94–1.03) | 0.47 | B |
| Weight-lowering | Female | 40 | 9,289 | 1.03 (0.98–1.07) | 0.22 | B |
| Non-lowering | Female | 48 | 10,504 | 1.03 (0.99–1.07) | 0.16 | B |
| Pre-dosing, all experiments | Male | 60 | 12,689 | 0.95 (0.91–0.99) | 0.018 | B |
| Pre-dosing, weight-lowering | Male | 30 | 7,458 | 0.89 (0.85–0.94) | 1.9 × 10<sup>−5</sup> | B |
| Pre-dosing, non-lowering | Male | 30 | 7,744 | 1.01 (0.96–1.07) | 0.64 | B |
| Published extenders at 12 months | Male | 21 | 5,807 | 0.90 (0.85–0.96) | 5.5 × 10<sup>−4</sup> | C |
| Published extenders at 12 months | Female | 11 | 3,068 | 0.99 (0.92–1.07) | 0.84 | C |
| 1.4-month weight | Male | 4 | 730 | 1.00 (0.85–1.17) | 0.97 | D |
| 1.4-month weight | Female | 2 | 553 | 1.03 (0.86–1.22) | 0.77 | D |

Interaction HRs are per +1 s.d. of relative weight; HR < 1 indicates that treatment became more favourable at higher weight. In joint mouse-clustered models, the male class contrast was HR 0.91 (95% CI 0.87–0.96; P = 2.2 × 10<sup>−4</sup>); in the pre-dosing subset it was HR 0.88 (P = 1.6 × 10<sup>−4</sup>). Primary and pre-dosing models used six-month weight, with follow-up from six months or treatment initiation, whichever was later. Mice can contribute to more than one experiment through shared controls; variance was clustered by mouse. Among published extenders, the male interaction was absent at 18 months (HR 1.01, 0.95–1.06; P = 0.81) and reversed at 24 months (HR 1.11, 1.03–1.18; P = 0.0031), paralleling the late-life reversal of the weight–mortality association in controls; at 6 months only two experiments qualified (HR 0.80, P = 0.0029). The 12-month estimate was similar with a 90-day lag (HR 0.91, 0.86–0.97; P = 0.0022) and after omitting each cohort in turn (HR 0.88–0.92; all P < 0.01). Female estimates showed no association at any landmark (all P > 0.05). Source files in the repository data/outputs folder: A, treatment\_moderation.csv, primary rows; B, the same file, weight\_class rows; C, the same file, landmark rows; D, developmental\_weights.csv, puberty\_moderation rows. Mouse-level inputs are provided in Supplementary Data 2.

Interventions also vary in the weight effect that they induce, so we analysed separately those that lowered body weight and those that did not. The relationship was particularly apparent among the 42 weight-lowering experiments, for which the corresponding difference was approximately 10% (HR 0.90, 0.86–0.94), while no clear weight dependence was detected among the 50 non-weight-lowering experiments (difference between classes, P = 2.2 × 10⁻⁴). Females showed no corresponding treatment effect moderation across 88 experiments (HR 1.03, 0.99–1.07; P = 0.11; Fig. 2a; Table 1). Examples of weight-lowering compounds include acarbose, 17α-estradiol, canagliflozin and rapamycin regimens (Supplementary Data 1).

We then repeated the analysis using only weights measured before dosing, to assess whether the pattern preceded treatment. The association remained across the 60 male experiments (HR 0.95, 0.91–0.99; P = 0.018; Table 1), with the same stronger weight dependence among weight-lowering interventions, whereas no significant effect was detected in females (HR 1.02, 0.99–1.07; P = 0.22; Table 1).

### Survival declines more steeply with weight in controls than in treated mice

To express the weight-dependent lifespan increase in days and percentage, we divided mice into pre-treatment weight quartiles across the 30 male weight-lowering experiments with weights recorded before dosing began, so that weight could not have been influenced by treatment. Quartiles were formed separately within each experiment and standardized so that each experiment and site contributed equally to every quartile.

Median lifespan gain increased from 55 days in the lightest quartile (890 to 945 days; +6.2%) to 136 days in the heaviest (673 to 809 days; +20.2%; Fig. 1b–d; Table 2), while the second and third quartiles showed increases of 8.0% and 12.3%. The greater increase in heavier mice arose mainly because control survival declined more steeply with weight: from the lightest to the heaviest quartile, median lifespan fell by 217 days in controls but by 136 days in treated animals (Fig. 1c,d). Nevertheless, treated mice in the heaviest quartile remained shorter-lived than untreated mice in the lightest quartile. Excess mortality in heavier controls began early, with 27.6% of male controls in the heaviest quartile dead by day 500 against 6.3% in the lightest.

**Table 2.**
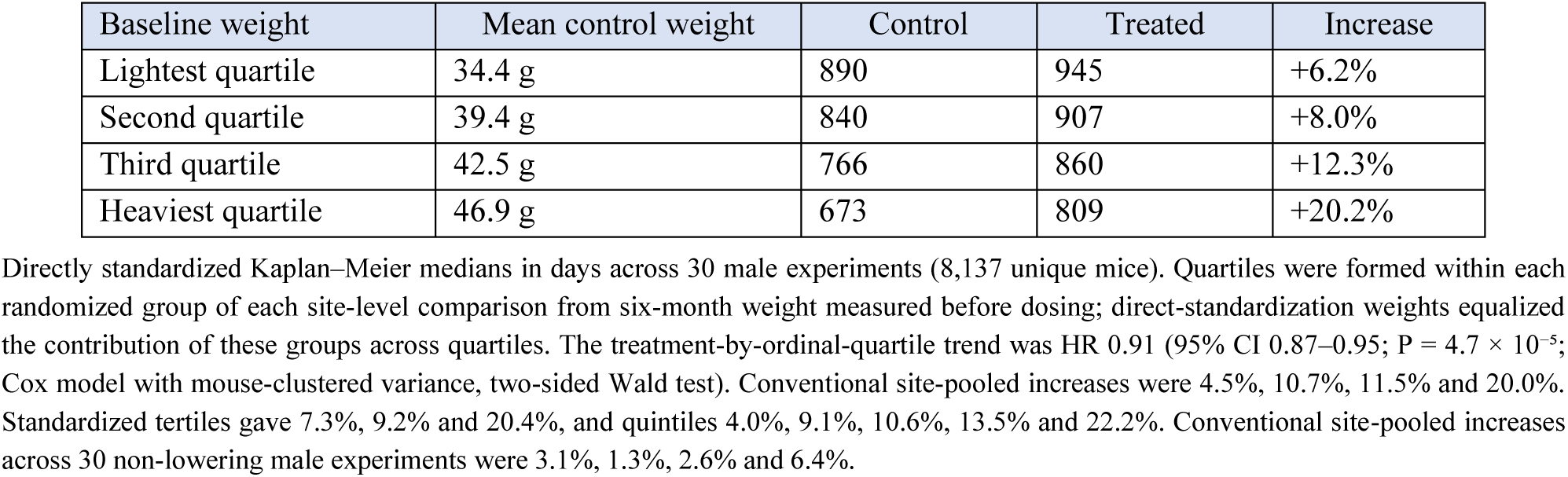
Median lifespan by quartile of pre-treatment body weight in male weight-lowering experiments.

| Baseline weight | Mean control weight | Control | Treated | Increase |
|---|---|---|---|---|
| Lightest quartile | 34.4 g | 890 | 945 | +6.2% |
| Second quartile | 39.4 g | 840 | 907 | +8.0% |
| Third quartile | 42.5 g | 766 | 860 | +12.3% |
| Heaviest quartile | 46.9 g | 673 | 809 | +20.2% |

Directly standardized Kaplan–Meier medians in days across 30 male experiments (8,137 unique mice). Quartiles were formed within each randomized group of each site-level comparison from six-month weight measured before dosing; direct-standardization weights equalized the contribution of these groups across quartiles. The treatment-by-ordinal-quartile trend was HR 0.91 (95% CI 0.87–0.95; P = 4.7 × 10<sup>−5</sup>; Cox model with mouse-clustered variance, two-sided Wald test). Conventional site-pooled increases were 4.5%, 10.7%, 11.5% and 20.0%. Standardized tertiles gave 7.3%, 9.2% and 20.4%, and quintiles 4.0%, 9.1%, 10.6%, 13.5% and 22.2%. Conventional site-pooled increases across 30 non-lowering male experiments were 3.1%, 1.3%, 2.6% and 6.4%.

In a Cox model, the treatment effect strengthened progressively from one weight quartile to the next (HR 0.91 per heavier quartile, 95% CI 0.87–0.95; P = 4.7 × 10⁻⁵; Fig. 2b). To check that the finding did not depend on median lifespan, we also compared mean survival accumulated from six months of age to 1,000 days. Treatment added 37.3 days in the lightest quartile and 88.9 days in the heaviest, a difference of 51.6 days (95% CI 26.7–76.9). The gradient was also retained using site-pooled medians, dividing animals into tertiles or quintiles, and applying each control group’s weight cut-offs to both randomized groups (Table 2). By contrast, the corresponding increases in 30 non-weight-lowering male experiments were smaller, and showed no consistent progression across quartiles: 3.1%, 1.3%, 2.6% and 6.4% (Fig. 2b; HR 1.01, 0.96–1.07; P = 0.64; Table 1).

Even the lowest-weight quartile had a mean control weight of 34.4 g, approximately 14% above the 30.3 g mean across 27 male laboratory strains measured at a similar age.^11^ Thus, the lowest-weight quartile represents a relative weight stratum rather than a metabolically lean or low-adiposity reference (Fig. 3a).

**Figure 3.**
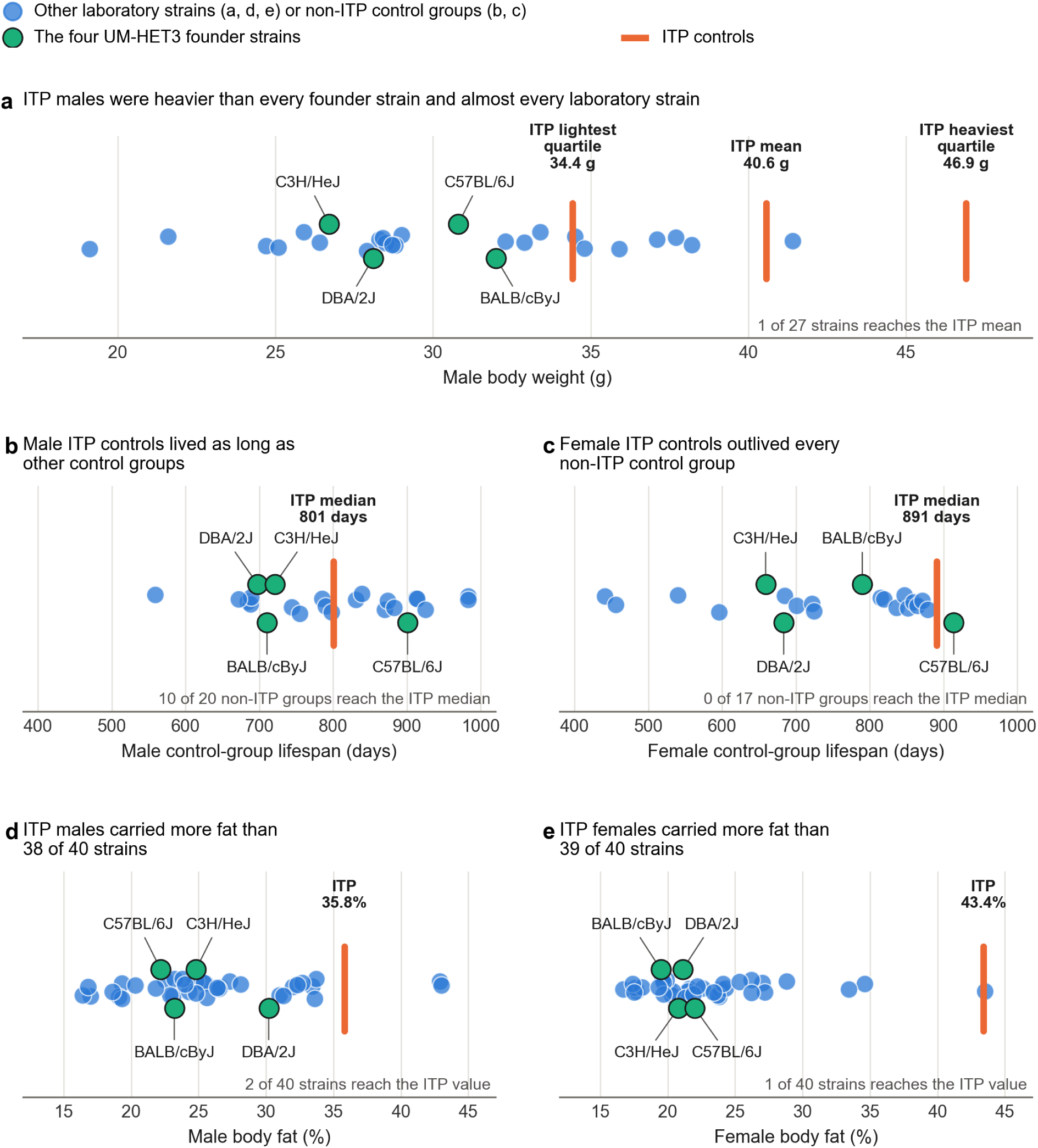
ITP controls are heavier and fatter than most laboratory strains, and ITP female controls outlive non-ITP female controls. Each dot is one laboratory strain (a, d, e) or one non-ITP control group (b, c), with vertical jitter; labelled dots are the four UM-HET3 founder strains and bars mark ITP controls. a, Male body weight, with average control weights in the lightest and heaviest pre-treatment quartiles and the overall six-month male control mean. b, Male control-group lifespan (18 ITP and 20 non-ITP groups). c, Female control-group lifespan (18 ITP and 17 non-ITP groups). d, Male body fat. e, Female body fat. Reference body weights come from the Reed1 survey^11^ (27 of 28 strains measured at 29–34 weeks); reference body fat comes from the 40-strain dual-energy X-ray absorptiometry survey^27^ (14–18 weeks). ITP weights were measured at six months, and ITP body fat was measured by whole-body MRI at nine months^28^. The scanned mice were distinct from those in our individual survival analyses. Founder-strain weights are reported in Extended Data Table 8. Control-group definitions and reported lifespan endpoints follow Extended Data Table 7; the bar shows the median across ITP control groups. Founder-strain median lifespans in b and c are from Yuan et al.^20,21^; these additional reference observations are excluded from the non-ITP control-group counts.

### The effect of body weight is consistent across interventions

To test whether the pooled gradient was confined to a small number of interventions, we grouped the 26 weight-lowering single-agent experiments by their 14 active compounds and compared lifespan increases across pre-treatment weight. The lightest animals retained only about one-quarter of the increase observed in the heaviest animals (5.0% versus 20.2%). The same ordering occurred for 13 of the 14 compounds, including acarbose, 17α-estradiol, canagliflozin and rapamycin (Fig. 4).^12–16^ The common treatment-by-weight interaction showed greater lifespan increase at higher weight (HR 0.90, 95% CI 0.85–0.95; P = 1.1 × 10⁻⁴). The pooled association was therefore not confined to a single compound or dosing schedule.

**Figure 4.**
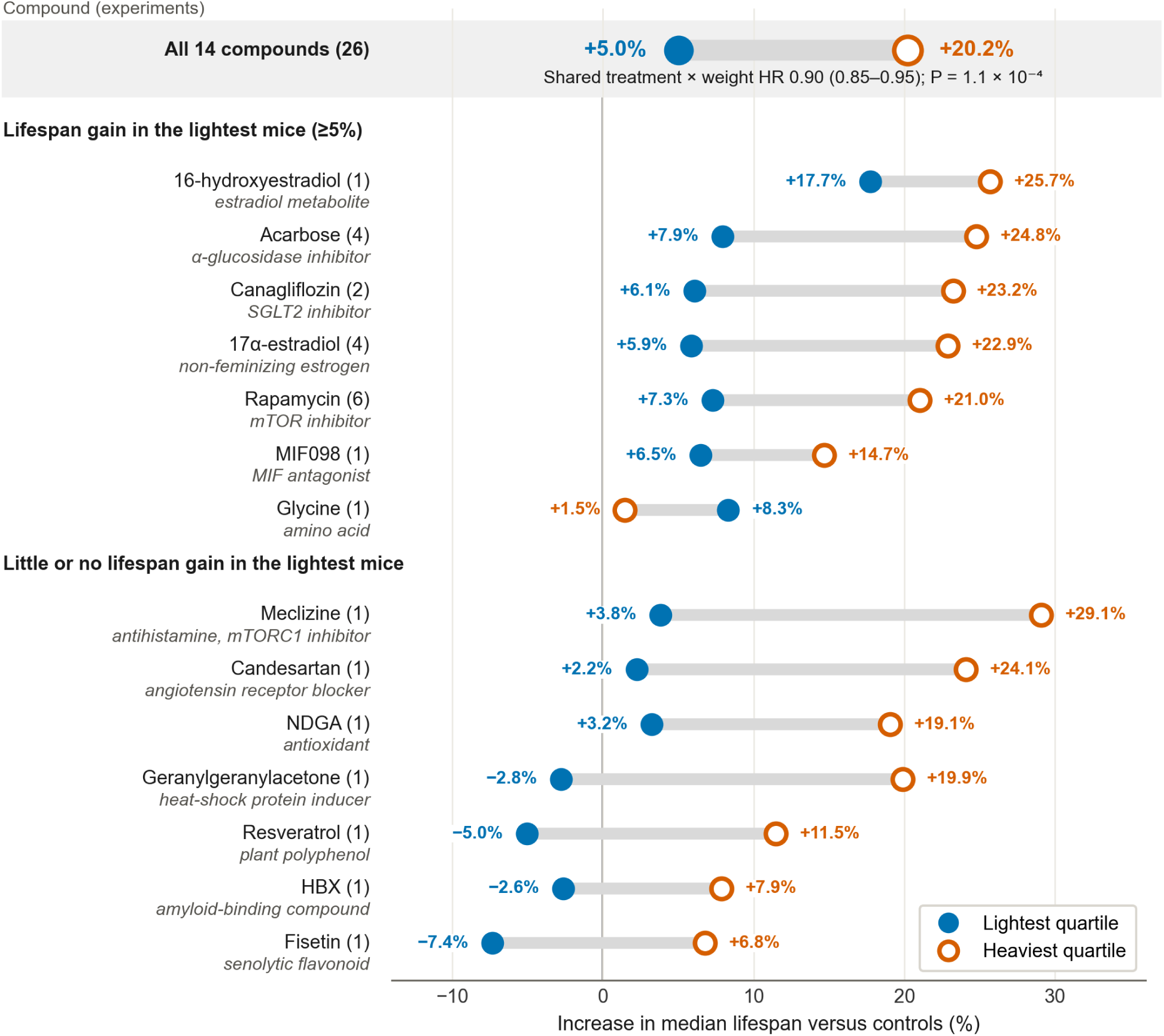
For 13 of 14 compounds, lifespan increase is larger in the heaviest than in the lightest quartile. Filled and open circles give the increase in median lifespan in the lightest and heaviest pre-treatment weight quartiles for each of the 14 single compounds tested in weight-lowering experiments in males, grouped by whether the lightest quartile had a lifespan gain of at least 5%. The drug class or mechanism of each compound is given beneath its name. The pooled single-agent set (top) combines all 26 experiments (7,238 mice). Values in parentheses give the number of experiments; HBX, 2-(2-hydroxyphenyl)benzoxazole. The common treatment-by-weight interaction HR (Cox model) is shown with its 95% confidence interval and two-sided Wald P value.

### Body weight predicts subsequent mortality

We next examined why heavier treated mice showed larger lifespan increases, starting with whether weight itself predicts death in untreated controls, as reported previously.^6,7^ It did, strongly in males and weakly in females: each standard deviation of six-month weight was associated with a 39% higher hazard of death in males and an 8% higher hazard in females (HR 1.39 and 1.08; Fig. 5a; Extended Data Table 1). The inverse association between weight and lifespan was nearly universal across cohorts and interventions, with heavier mice living shorter lives in 108 of 109 male and 87 of 105 female groups. The difference in magnitude persisted at twelve months (male HR 1.36; female HR 1.10; Fig. 5a) and weakened by eighteen months. By 24 months the male association had reversed, consistent with selective survival and weight loss before death. The six-month gradient was unchanged after adjustment for maternal age and parity, both estimates shifting by less than 0.01.

**Figure 5.**
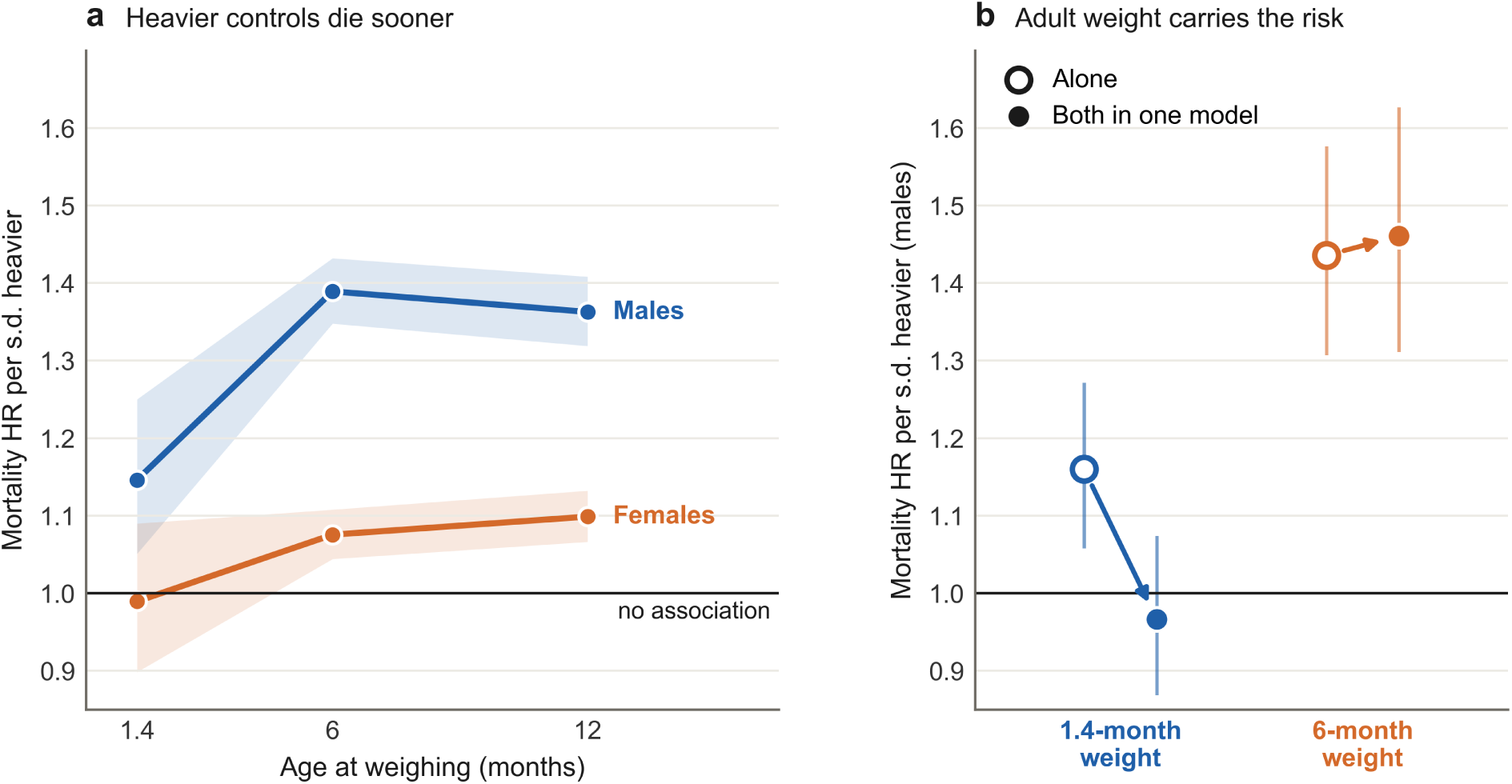
Adult weight, not early-life weight, independently predicts mortality in controls. a, Subsequent mortality HRs (Cox models) for weight measured at 1.4, 6 and 12 months in untreated controls. b, 1.4- and 6-month weight, each modelled alone (open circles) or both in one model (filled circles), among 539 male controls with both measurements. Shading and error bars show 95% confidence intervals; sample sizes are given in Extended Data Table 1.

### The effect of weight emerged with acquired adult weight

Higher adult weight may reflect a larger body frame, greater adiposity, or both. To determine the roles of early life intrinsic body size and weight gained subsequently, we examined weight at puberty (approximately 1.4 months of age), available in three cohorts. Early-life weight did not modify the treatment effect in either sex (males: HR 1.00, 0.85–1.17; P = 0.97; females: HR 1.03, 0.86–1.22; P = 0.77; Table 1), and pubertal male weight did not predict mortality once six-month weight was taken into account (HR 0.97 per s.d., 0.87–1.07; P = 0.52), whereas six-month weight remained strongly predictive (HR 1.46, 1.31–1.62; P = 2.7 × 10⁻¹²; Fig. 5b).

To distinguish weight already attained at adulthood from subsequent weight gain, we also examined changes between six and twelve months among mice surviving beyond twelve months. In controls and interventions initiated before twelve months, greater gain predicted higher subsequent male mortality even after adjustment for six-month weight (HR 1.10, 95% CI 1.07–1.12; 14,118 mice). The corresponding association was smaller in females (HR 1.04, 1.03–1.06; 13,167 mice). The male association remained when controls were excluded and when the analysis instead included late-start interventions, for which the six-to-twelve-month gain preceded dosing. Together with the early-life analysis, these findings support attained adult weight and subsequent gain as the relevant risk phenotype, rather than constitutional body size alone.

### Heavier control groups yield larger treatment-effect estimates

We next asked whether apparent intervention efficacy also differed between experiments according to the average weight of the control group. We split each experiment by site, giving 534 site-level comparisons, and grouped them by their own controls’ six-month weight. In males, pooled hazard reduction approximately doubled from 10.9% against the lightest comparator tertile to 21.5% against the heaviest, while mean median-lifespan increase rose from 4.0% to 8.3% (Table 3). The association remained in within-cohort comparisons accounting for shared controls (HR 0.96 per s.d., 95% CI 0.931–0.997; P = 0.034). Female estimates showed the same pattern, with hazard reduction rising from 5.3% to 12.0% and median-lifespan increase from 1.9% to 2.8%, but the corresponding within-cohort estimate was not statistically significant (P = 0.27).

**Table 3.**
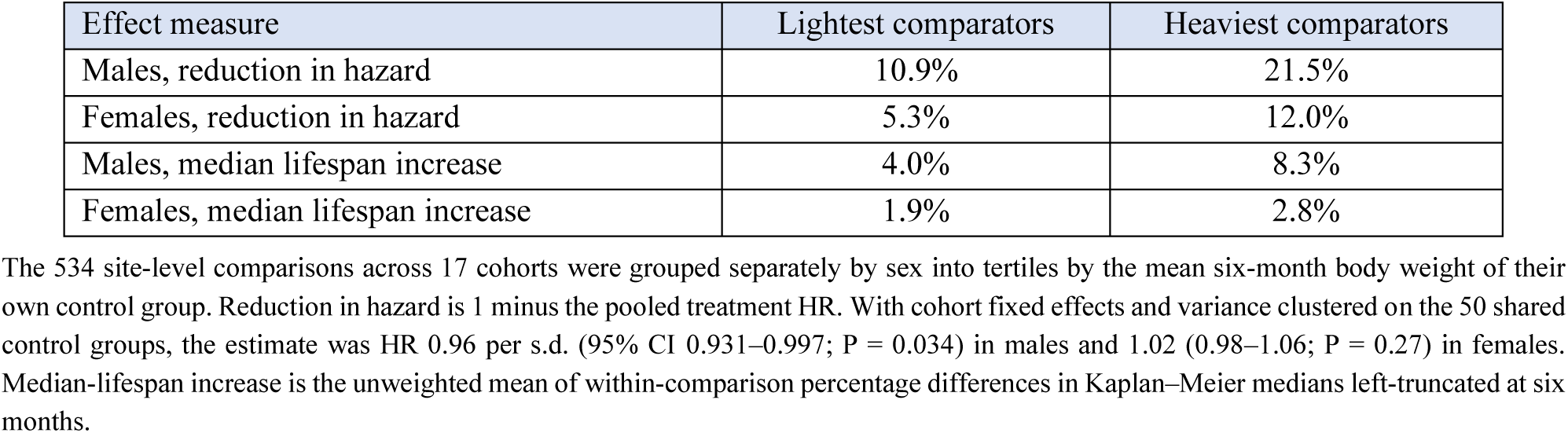
Size of the treatment effect against the lowest-weight and highest-weight comparator groups.

| Effect measure | Lightest comparators | Heaviest comparators |
|---|---|---|
| Males, reduction in hazard | 10.9% | 21.5% |
| Females, reduction in hazard | 5.3% | 12.0% |
| Males, median lifespan increase | 4.0% | 8.3% |
| Females, median lifespan increase | 1.9% | 2.8% |

The 534 site-level comparisons across 17 cohorts were grouped separately by sex into tertiles by the mean six-month body weight of their own control group. Reduction in hazard is 1 minus the pooled treatment HR. With cohort fixed effects and variance clustered on the 50 shared control groups, the estimate was HR 0.96 per s.d. (95% CI 0.931–0.997; P = 0.034) in males and 1.02 (0.98–1.06; P = 0.27) in females. Median-lifespan increase is the unweighted mean of within-comparison percentage differences in Kaplan–Meier medians left-truncated at six months.

### Larger increases in lifespan in experiments with greater weight loss in the intervention group

We examined whether interventions producing larger average weight reductions also produced larger lifespan increases, as previously reported.^4^ In males, each 1% greater weight reduction was associated with 0.71 percentage points greater lifespan increase (95% CI 0.32–1.10; P = 0.0016; 83 experiments), and the association persisted within cohorts (0.65, 0.20–1.10; P = 0.0078). Mean lifespan increase rose from 2.7% in the quartile with the least weight loss to 10.9% in the greatest-loss quartile (P = 0.018). The corresponding female within-cohort association was weaker (0.17, −0.15 to 0.50; P = 0.28; Extended Data Table 2). The relationship held after adjustment for comparator weight. Across 246 site-level comparisons for males, the treatment-associated weight-change coefficient was essentially unchanged (0.022 versus 0.021 log hazard ratio per 1% higher weight; adjusted 95% CI 0.014–0.029; P = 1.2 × 10⁻⁶), and the predictors were weakly correlated (Spearman ρ = −0.12). In females, the adjusted coefficient was smaller but also positive (0.011, 0.004–0.019; P = 0.0043).

### The attenuated weight dependence in females is a feature of the ITP

To determine whether the weaker female treatment-by-weight association was confined to one ITP site, we conducted site-specific analyses. Female estimates were close to null at all three sites, with no evidence of between-site heterogeneity (P = 0.62). To determine whether the attenuated effect in females was a feature of other studies, we made a cross-study comparison for experiments that both extended lifespan and lowered weight. Lifespan increase scaled more strongly with weight reduction in ITP males than females (1.00 versus 0.43 percentage points; P = 0.017 for the difference) as previously reported,^4^ whereas outside ITP the slopes were similar in the two sexes (0.70 in males and 0.86 in females; P = 0.46). This sex-by-source difference was significant (P = 0.018), including when only experiments with at least 5% weight loss were analysed (Extended Data Table 3).

An independent female Diversity Outbred cohort provided an individual-level external contrast.^17^ Greater pre-intervention weight predicted subsequent mortality under *ad libitum* feeding, whereas no clear gradient was detected under caloric restriction. Lifespan increase rose from 96 days (10.9%) in the lightest quartile to 232 days (32.5%) in the heaviest. Across those quartiles, *ad libitum* median survival fell from 877 to 714 days, while restricted median survival changed modestly from 973 to 946 days. The ordering resembled the male ITP pattern, although the treatment-by-quartile estimate was not significant (HR 0.90, 95% CI 0.76–1.05; P = 0.18; Extended Data Table 4). A second genetically heterogeneous cohort from an olfactory-exposure lifespan study showed the same pattern among females,^18^ with greater six-month weight predicting subsequent mortality (HR 1.52 per s.d., 95% CI 1.14–2.02; P = 0.0042; 59 control mice), whereas six-week weight did not once adult weight was included (Extended Data Table 5). The weaker female weight dependence therefore appeared to be concentrated within the ITP.

### ITP females have low early mortality and high control survival

The weaker weight dependence of lifespan in ITP females than in males could reflect unusually long-lived controls. In pooled untreated controls from all 17 ITP cohorts, Kaplan–Meier median survival showed an 86-day female advantage over males, present in every cohort (range +52 to +135 days) and consistent with previous sex-specific survival analyses of six of these cohorts.^19^ A large female survival advantage is, however, not a general feature of laboratory mice. Across 31 inbred strains, the median female–male difference in median lifespan was 9 days, compared with 86 days in ITP, and only 2 of 31 strains exceeded the ITP female median lifespan of 886 days.^20,21^

By day 500, Kaplan–Meier mortality was 2.4% in ITP females and 16.0% in males. By comparison, 12.2% of *ad libitum* female Diversity Outbred controls had died by the same age, approximately four times the corresponding ITP female value (3.0%).^17^ Independent UM-HET3 controls differed by only 5 days in mean lifespan (836 versus 831 days),^22^ whereas an independent C57BL/6J cohort maintained on the same 5LG6 chow showed a 61-day female advantage in median survival (865 versus 804 days).^23^ Together, these comparisons place ITP females within a control population with substantially less early and midlife mortality available for interventions to prevent (Extended Data Table 6).

ITP female control groups also occupied the upper end of the sampled control-survival distribution. Median control survival was 891 days across 18 ITP female control groups and 815 days across 17 non-ITP female control groups, and no sampled non-ITP control group reached the ITP median (P = 2.5 × 10⁻⁶). The contrast persisted when each study contributed a single sex-specific median (893 versus 724 days; 13 and 15 studies; P = 1.5 × 10⁻⁵) and when analysis was restricted to median-lifespan endpoints (891 versus 847 days; 0 of 9 non-ITP groups reached the ITP median; P = 2.9 × 10⁻⁴). Restricting the comparison to studies published from 2000 onward gave the same pattern (891 versus 842 days; 0 of 12; P = 3.6 × 10⁻⁵). Male control survival did not differ under any of these specifications (all P > 0.1). Thus, the ITP female lifespans lay above the sampled non-ITP range, whereas the male distributions overlapped (Fig. 3b,c; Extended Data Table 7).

### Population architecture may contribute to the ITP-specific pattern

Some features of the ITP setting could bear on the weaker association between weight and survival in females. UM-HET3 mice descend from four inbred founders, while other cohorts in the sampled studies, such as Diversity Outbred mice, derive from eight founders, including three wild-derived strains and others susceptible to obesity or autoimmune diabetes.^17,24–26^ This broader genetic architecture may encompass a wider range of metabolic susceptibility, allowing adult weight to mark accumulated risk differently across populations.

ITP controls were also substantially heavier and more adipose than their founder strains. The four founders averaged 29.4 g at approximately seven months in an independent Mouse Phenome Database survey (Extended Data Table 8), whereas ITP male controls averaged 40.6 g at six months, 38.0% heavier despite being measured one month earlier. Even the lightest ITP quartile exceeded every founder strain’s mean weight. Although heterosis may contribute to larger body size, ITP controls also had higher percentage body fat than their founders (35.8% versus 25.1% in males and 43.4% versus 20.9% in females). Only two of the 40 surveyed strains reached the male ITP adiposity value and one the female value (Fig. 3d,e).^11,27,28^

Continuous access to food within sedentary laboratory housing may contribute to this metabolic state.^1^ In UM-HET3 mice, controlled food availability lowered eight-month weight from 45 to 29 g in males and from 35 to 22 g in females while increasing mean lifespan from 831 to 1,101 days (+32%) and from 836 to 1,169 days (+40%), respectively.^22^ Together with the stronger weight–mortality associations observed in other female cohorts, these findings suggest that the attenuated weight–mortality association in ITP females may be population- or husbandry-specific rather than sex-specific.

## Discussion

We have found that the magnitude of intervention-generated increase in lifespan in male mice depends in part on attained adult weight. Treatment-by-weight interaction was detected across 92 experiments and concentrated among interventions that lowered weight. This dependence arose mainly because control survival declined with weight more steeply than treated survival: even with treatment, animals in the heaviest quartile died earlier than untreated animals in the lightest (809 versus 890 days). The same dependence was evident at the compound level, where lifespan extension was approximately four times as large in the heaviest weight quartile as in the lightest, with the same ordering for 13 of 14 compounds. Together, these patterns appear consistent with partial rescue of weight-associated risk rather than elevation of a lifespan ceiling.

The effects on lifespan appear to reflect weight acquired during adulthood rather than constitutional body size. Early-life weight predicted mortality in isolation but was not independently associated once adult weight was included, whereas six-month weight remained strongly predictive and subsequent adult gain added risk. The relevant treatment variable is therefore better described as attained adult weight rather than as body size established early in life. Complementary experiment-level analyses showed that greater treatment-associated weight reduction accompanied larger male lifespan increases within cohorts, and its association with treatment hazard ratios persisted after adjustment for comparator weight.

The attenuated weight effect in females was concentrated in ITP rather than shared across the sampled mouse intervention studies. Within ITP, male control mortality rose steeply with adult weight, whereas the female gradient was smaller. Outside ITP, lifespan increase scaled similarly with weight reduction in both sexes. Control survival suggests one contributor: ITP females had particularly low early and midlife mortality, and none of the sampled non-ITP female control groups reached the median control survival of ITP females, whereas the corresponding male distributions overlapped. Where female *ad libitum* controls carried more early mortality, as in Diversity Outbred mice, restriction benefit rose from 10.9% in the lightest weight quartile to 32.5% in the heaviest.^17^ This pattern appears more consistent with an ITP-specific population or husbandry effect than with a broadly attenuated association in females.

The weight dependence of treatment effects also bears on what constitutes an appropriate comparator. In humans, body-weight ranges are defined in part by their association with morbidity and mortality, with the lowest all-cause mortality falling within the normal BMI range and risk rising below and above it.^29,30^ Mouse body weight cannot be mapped directly onto human BMI, but the underlying risk-based logic still applies: a comparator state in which greater weight is accompanied by substantially higher mortality may not represent an equivalent healthy baseline. Among untreated male ITP controls, greater six-month weight was strongly associated with subsequent mortality (HR 1.39 per s.d., 1.35–1.43) and, across the matched pre-treatment quartiles, mean control weight rose from 34.4 to 46.9 g, while standardized control median lifespan fell monotonically from 890 to 673 days. The heavier mice therefore appear to represent progressively higher-risk physiological states, so that their larger treatment effects may partly reflect rescue of weight-associated mortality rather than longevity gain from treatment. A meta-analysis likewise showed lower mortality associated with intentional weight loss among people with obesity-related risk factors, alongside higher mortality among those healthy at baseline,^31^ suggesting that the association between weight loss and survival depends on baseline physiological state. While greater benefit in higher-risk animals could be valuable for individuals carrying similar risk, the magnitude of lifespan extension may not generalize to metabolically healthier populations. Differences in baseline metabolic risk between experimental controls and clinical populations may contribute to the difficulty of translating preclinical results.^1^

Only the lightest quartile approached the approximately 900-day reference proposed for adequately long-lived laboratory controls,^5^ and in the lightest quintile, which reached it, the increase was lower still (4.0% versus 6.2%). Because mortality kept falling with lower weight across the whole measured range, without the rise at low weight normally seen for BMI in humans,^29,30^ these lighter mice may represent the closest available approximation to a lower-risk comparator rather than an established metabolically optimal state. They nonetheless remained fed *ad libitum* and heavier than UM-HET3 males under controlled feeding (34.4 g versus 29 g), a regimen that extended mean lifespan by 32%,^22^ so whether their observed lifespan increases would be maintained against a metabolically healthier comparator remains unclear. The compound-level results (Fig. 4) suggest which interventions might retain benefit against healthier controls. In the lightest quartile, acarbose, rapamycin, canagliflozin and 17α-estradiol still increased median lifespan by 6–8%, against 21–25% in the heaviest. All four act on nutrient-sensing or glucose-handling pathways (carbohydrate digestion, urinary glucose loss, mTOR and hypothalamic feeding circuits),^12,14,28,32^ suggesting actions beyond weight loss. Others, including resveratrol and fisetin, showed no increase in the lightest quartile, although most compounds were tested in only one experiment.

Non-human-primate studies provide a parallel comparator contrast. In the Wisconsin rhesus study, where controls approximated *ad libitum* feeding, caloric restriction improved all-cause survival,^33,34^ whereas no significant survival extension was detected at the NIA, where controls were portioned to avoid overfeeding and were often as lean as Wisconsin restricted animals.^34,35^ Although the studies differed in age at initiation, diet and ancestry, the divergence is consistent with the apparent benefit of restriction depending partly on the baseline metabolic state of the comparator. This interpretation accords with longstanding concerns that continuously fed, sedentary laboratory rodents can develop excess weight, impaired glucose regulation and premature mortality, and that interventions effective in this state may reproduce less strongly in active, normal-weight populations.^1^

When *ad libitum* husbandry contributes to metabolic dysfunction, treatments that reduce food intake through altered appetite or palatability may improve disease outcomes partly by correcting that state rather than through disease-specific pharmacology.^1^ This question is becoming more relevant as appetite-suppressing drugs enter aging research. Late-life semaglutide reduced food intake by about 24% and extended median lifespan by about 12% in old, *ad libitum*-fed female mice.^36^ While an intake-matched caloric-restriction comparison showed functional benefits, how much of the lifespan increase would remain against a leaner, portion-fed comparator is untested. The implications may extend beyond lifespan studies. In regulatory toxicology, body weight has long been known to predict the incidence of certain spontaneous tumours, and treatment-related weight reductions could lower tumour incidence or mask carcinogenic effects.^37,38^

These findings suggest several priorities for future experiments. Most published lifespan studies do not report effects across baseline weight and metabolic-risk strata, limiting how directly control-state dependence can be assessed. Reporting individual body weights alongside survival would show whether benefit is concentrated in heavier animals or persists across the weight range. Leading interventions could also be tested against both metabolically challenged and healthier comparators, for example under *ad libitum* and controlled meal-feeding,^39^ to determine whether benefits depend on continuous food access. Pair-fed or weight-matched controls could further distinguish effects reproduced by altered energy balance from effects beyond it, particularly for compounds whose effects on intake can mimic dietary restriction.^40^ Finally, replication across genetically distinct populations would help establish the generalizability of these findings across different metabolic susceptibilities.

These analyses concern variation within *ad libitum*-fed populations and do not determine how far the *ad libitum* state itself is displaced from a metabolically healthy baseline. Virtually all sampled lifespan experiments compared treatment against *ad libitum*-fed controls, whereas survival effects against metabolically healthy, non-*ad libitum* comparators remain largely untested. The mouse data suggest that efficacy may not be fully maintained as baseline metabolic risk falls.

In conclusion, attained adult weight modified male geroprotective efficacy in the ITP, with larger effects in heavier animals and for weight-lowering interventions. Although these findings warrant further investigation across additional mouse strains and under controlled-feeding conditions, large effects against heavy, short-lived controls may partly reflect rescue conditional on comparator state, and their magnitude may not generalize to metabolically healthier comparators.

## Methods

### Data sources and harmonization

Individual body weights were obtained from the Mouse Phenome Database ITP cohort projects (ITP2004–ITP2021) and GeneNetwork HET3-ITP, and survival records from Mouse Phenome Database ITP1, across 17 cohorts (Supplementary Data 1), yielding 36,973 unique animals, of which 35,476 had at least one recorded adult weight (Supplementary Data 2).^9,10,41^ Weights were linked by cohort and normalized mouse identifier, with sex, site, treatment and overlapping measurements checked. Treatment-group discrepancies were resolved using survival records after cross-checking the linked weights. Adult weights were withheld for two mice with unresolved sex discrepancies; their survival records were retained. Weight records without reliable survival linkage were excluded. Duplicate 2020 survival records were removed, and weights outside 10–100 g were treated as recording errors. The primary treatment-moderation model retained 33,780 unique mice after experiment and age eligibility. Death and censoring definitions followed the source files; controls reused across intervention comparisons were retained with variance clustered by mouse.

### Treatment-moderation model

The primary analysis included every intervention experiment with treated and control animals, survival beyond the start of follow-up, a usable six-month weight and at least 40 deaths to limit unstable treatment-by-weight estimates. Six-month weight provided a common adult measurement across experiments. Follow-up began at six months or at treatment initiation, whichever was later, with concurrent controls entering at the same age, so that deaths before dosing did not enter the comparison. Weight was z-standardized within treatment group × site × sex, so that the treatment-by-weight interaction reflected differences in relative weight within each group rather than weight differences between treated and control animals. Cox models were left-truncated at the start of follow-up, baseline hazards were stratified by site-level comparison (one experiment at one site, in one sex), treatment and weight main effects were experiment-specific, the treatment-by-weight coefficient was shared, and robust variance was clustered by mouse. Consistency across sites was tested by allowing site-specific treatment, weight and treatment-by-weight terms and comparing the treatment-by-weight terms with a robust Wald test. HR < 1 indicates a more favourable treatment effect at higher within-experiment weight. Interventions were classified as weight-lowering when treated animals averaged more than 1 g (about 2.5% of mean six-month control weight) below their own controls across available post-dosing adult measurements, weighted by treated sample size. This classifier used weight response rather than survival outcome. Differences between the weight-lowering and non-lowering classes were tested in joint Cox models containing treatment-by-weight-by-class terms with mouse-clustered variance. Because six-month weight followed the start of treatment in experiments dosed by six months, the class analysis was repeated using only weights measured before dosing. The matched-quartile, compound and comparator analyses instead measured survival from six months, giving a common origin across starting ages. A secondary analysis was restricted to intervention–sex combinations reported to extend lifespan in the corresponding ITP publication (Supplementary Data 1). Models were fitted at 6, 12, 18 and 24 months, with emphasis on twelve months as the best compromise between established adult weight and limited mortality selection. A 90-day lag excluded deaths shortly after each landmark. Leave-one-cohort-out analyses assessed sensitivity to individual cohorts.

### Comparator-state analyses

For the matched-quartile analysis, 30 male weight-lowering experiments with a six-month measurement before dosing were included (Supplementary Data 3). To prevent differences in cohort, site, intervention or randomized-group composition from driving the gradient, weight quartiles were assigned within each randomized group of each site-level comparison. Direct-standardization weights then gave every randomized group of each eligible site-level comparison the same relative contribution in each quartile. Weighted Kaplan–Meier medians were calculated for treated and control animals, and proportional increase was 100 × (treated median − control median)/control median. A left-truncated Cox model stratified by site-level comparison, clustered by mouse and allowing experiment-specific treatment effects estimated the shared treatment-by-ordinal-quartile trend. Restricted mean survival from six months to age 1,000 days used the same standardization, with 95% percentile intervals from 2,000 mouse-level Bayesian bootstrap replicates in which each mouse received the same bootstrap multiplier wherever its record was reused. Sensitivity analyses comprised conventional quartiles, with weight cut-offs computed within site across experiments and randomized groups; tertiles and quintiles formed in the same way as the primary quartiles; and quartiles defined by the control group’s cut-offs in each site-level comparison, applied to both randomized groups (88 of 89 comparisons represented in every quartile). Conventional quartiles also summarized the 30 non-lowering male experiments with pre-dosing weights. Day-500 mortality was estimated from the standardized control survival curves.

For the comparator-state analysis, treated mice in each site-level comparison were compared with their own contemporaneous control group. Comparisons required at least 25 treated and 25 control animals, survival beyond the six-month landmark and at least 30 deaths across the two groups. Within each eligible comparison, a Cox proportional-hazards model, left-truncated at six months, estimated the treatment log hazard ratio. Each animal appeared once within a comparison-level model; control animals could, however, recur across comparisons when one control group was shared by several intervention experiments. Comparison-level log hazard ratios were related to the mean six-month weight of the corresponding control group by inverse-variance-weighted regression. Control-group weight, rather than control survival, indexed comparator state because it is measured independently of the survival outcome. The model included cohort fixed effects and control-group weight standardized separately within sex, with cluster-robust variance at the cohort × site control-group level (50 clusters) because controls were shared across comparisons. Kaplan–Meier medians were estimated within each comparison after left truncation at six months. Percentage median-lifespan increase was calculated within each comparison as 100 × (treated median − control median) / control median and summarized as the unweighted mean across comparisons in each sex-specific comparator-weight tertile. Comparator-weight tertiles were used for descriptive presentation only; inferential analyses treated control-group weight continuously.

For the single-compound analysis, the 30-experiment set was collapsed by active compound after excluding four multicomponent experiments. All 14 remaining compounds were retained without selection by lifespan effect, and the same direct-standardization weights were used to estimate overall, lowest-quartile and highest-quartile Kaplan–Meier medians. Across the 26 single-agent experiments, the common-interaction model retained experiment-specific treatment and weight main effects and a shared treatment-by-weight term, matching the primary specification.

Kaplan–Meier median survival and mortality by day 500 were calculated in untreated controls from all 17 available ITP cohorts, censoring animals removed for fighting, accident or experimental procedures. For external context, sex-specific control survival was extracted from an independent UM-HET3 study and from an independent C57BL/6J study using the same 5LG6 chow. These rows were selected to match ancestry and diet separately. The Diversity Outbred source flags censored animals but not the reason for removal, so its day-500 estimate counted every animal as an event; the ITP female estimate was recalculated on the same basis (3.0%) for comparison.

### 1.4-month weight and adult-gain models

GeneNetwork trait ITP_50601 supplied body weight at approximately 42 days of age (1.4 months) in three cohorts.^10^ Early-life and six-month weights were z-standardized within cohort × site and entered separately and jointly in Cox models left-truncated at six months. The joint male-control model included 539 mice with both measurements. Treatment moderation at 1.4 months used published-extender experiments with at least 15 treated and 15 control mice, with follow-up from treatment start; weights were standardized against controls within cohort, site and sex. The primary trajectory analysis included controls and interventions initiated before twelve months across all seventeen cohorts, requiring six- and twelve-month weights and survival beyond day 365. Percentage gain was centred within each randomized group at each site and scaled by the pooled within-group s.d. separately by sex; groups required at least twelve deaths. Models stratified baseline hazards by group, used Efron ties and group-clustered robust variance, and estimated gain before and after adjustment for similarly centred six-month weight. The pre-treatment sensitivity instead included controls and interventions beginning at or after twelve months; it shared controls with the primary set but introduced late-start treated animals absent from that set. Maternal-age and parity models used the subset with both covariates. A treated-only sensitivity retained interventions initiated before twelve months and otherwise used the same group stratification, centering, scaling and six-month-weight adjustment.

### Sex-specific weight–mortality models

Among untreated controls, separate sex-specific Cox models related weight at 6, 12, 18 and 24 months to subsequent mortality. Weight was standardized within cohort × site × sex and baseline hazards were stratified by cohort and site. As a descriptive consistency analysis, site-specific Spearman correlations between six-month weight and age at natural death were calculated within randomized groups containing at least twelve animals, Fisher-z transformed and pooled across sites with n − 3 weights (Supplementary Data 4).

### External population context

UM-HET3 and Diversity Outbred founder architecture was summarized from published descriptions.^9,24,42^ UM-HET3 percentage fat came from the published nine-month MRI substudy.^28^ Founder-strain weights and ages were taken from Mouse Phenome Database Reed1 measures 32603 and 32680; housing and diet were taken from the corresponding project animal documentation.^11^ Mean six-month ITP weight was taken over all 4,983 male controls alive and weighed at that landmark. Strain percentage-fat values were taken from the 40-strain dual-energy X-ray absorptiometry survey; founder means were the unweighted means of the four UM-HET3 founder strains in that survey (Supplementary Data 5).

### Within-ITP experiment-level weight-change analysis

Within ITP, each treated group was compared with its concurrent cohort- and sex-matched control group. Percentage weight change used mean eighteen-month body weight, and percentage lifespan change used median age at natural death. Eligible experiments began before eighteen months so that weight was measured after dosing began and required at least eight treated and eight control observations for each measure, yielding 163 contrasts without selection on weight or lifespan change (Supplementary Data 6). Sex-specific ordinary least-squares models used cohort-clustered robust variance. Within-cohort slopes were estimated after centring both changes within cohort. Weight-change quartiles were formed separately within cohort and sex among cohorts with at least four eligible experiments, yielding 74 male and 68 female contrasts. An ordinal quartile term tested the trend, reported toward greater weight loss.

For comparator adjustment, experiment-level weight changes were joined to six-month-landmark treatment log hazard ratios by cohort, sex and intervention, yielding 246 male and 237 female site-level comparisons. Inverse-variance-weighted models included weight change alone or together with mean six-month control weight, with robust variance clustered by shared cohort × site control group. Weight-change coefficients are reported per 1% lower weight for lifespan increase and per 1% higher weight for log hazard ratio.

### External individual-level cohorts

Datasets for an external, individual-level test were sought among the studies in the cross-study corpus and in public repositories and journal source-data files of mouse lifespan studies (searched October 2026). The same eligibility criteria were applied to all 58 non-ITP studies in the cross-study corpus. A dataset was eligible if it compared a randomized intervention with *ad libitum*-fed controls, deposited body weight and survival for individual mice linked by identifier, used a population genetically heterogeneous within experimental groups, and recorded linked adult weight by twelve months of age to limit the influence of late-life disease-associated weight loss. Two cohorts met these criteria: the female Diversity Outbred cohort^17^ and mixed B6;129 mice from an olfactory-exposure lifespan study.^18^ The datasets assessed and the reasons for exclusion are listed in Supplementary Data 7. For the olfactory-exposure cohort, Cox models of mortality from six months were stratified by genotype, with additional exposure strata in pooled models, and used model-based variance. Treatment-by-weight interactions at twelve weeks and six and twelve months were fitted separately by sex and exposure, stratified by genotype (Supplementary Data 8).

The Dietary Restriction in Diversity Outbred mice (DRiDO) female, caloric-restriction groups were used as external individual-level contrasts rather than pooled with ITP intervention experiments.^17^ For the individual-level DRiDO analysis, female *ad libitum* and caloric-restriction groups were limited to mice with body weight measured within 45 days of the 180-day intervention landmark and surviving beyond that landmark. Pre-intervention weight quartiles were formed separately within the *ad libitum* and pooled caloric-restriction groups. The ordinal Cox model was left-truncated at 180 days and included separate main effects for each restriction level with a shared pooled-restriction-by-quartile term. Per-standard-deviation weight–mortality gradients were fitted separately within the *ad libitum* and pooled restricted groups.

### Cross-study intervention analysis

The cross-study corpus used mouse experiments recorded in DrugAge Build 5, supplemented by all subsequent original ITP lifespan reports published through October 2026. It contained 284 experiments from 69 studies reporting both treatment-associated weight change and lifespan change (Supplementary Data 9). One pooled-sex experiment contributed no sex-specific comparison, leaving 283 sex-specific intervention experiments from 68 studies for analysis. The responder analysis was restricted to 102 experiments in which treatment decreased body weight and increased lifespan, to characterize the magnitude of lifespan increase per unit weight reduction among interventions showing both responses. Experiments were classified as ITP or non-ITP interventions. Within each source-by-sex stratum, percentage lifespan increase was regressed through the origin on percentage body-weight reduction, such that the slope represents lifespan increase per 1% weight reduction, with standard errors clustered on study. Sex and source differences were tested by interactions with the weight term in the same clustered model, and the analysis was repeated after requiring at least 5% weight reduction. Weight and lifespan were obtained from DrugAge^43^ and the subsequent ITP reports, including mean and median lifespan measures and directly reported or figure-estimated weights.^4^ Source-supported post-dosing weights filled two missing DrugAge entries; source corrections are identified in Supplementary Data 9. Subsequent ITP additions required independent source support and a post-dosing weight measure. Because these comparisons are drawn between studies rather than within randomized experiments, study-level differences in husbandry, diet and survival ascertainment may confound these cross-study associations, which are therefore interpreted descriptively.

The control-lifespan comparison included 221 experiments from 40 DrugAge-listed or ITP studies reporting absolute control-group lifespan, independently of whether a numerical weight change was available: 80 ITP and 19 non-ITP female experiments, 90 ITP and 28 non-ITP male experiments, and four pooled-sex experiments excluded from sex-specific comparisons. ITP status required membership in an Interventions Testing Program cohort; UM-HET3 studies conducted outside the programme were classified as non-ITP. Because ITP publications commonly report multiple intervention experiments against one shared control group, experiments sharing a study and reported control lifespan were treated as one control group, leaving 18 ITP and 17 non-ITP female groups and 18 ITP and 20 non-ITP male groups. Control survival was compared within sex using two-sided Mann–Whitney tests. Because the studies spanned 1958 to 2026 and differed in the lifespan endpoint reported, sensitivity analyses allowed each study to contribute one sex-specific median, retained only median-lifespan reports, or restricted the comparison to studies published from 2000 onward (Extended Data Table 7).

### Statistical analysis

Cox proportional-hazards models used Efron handling of ties and normal-reference Wald tests. Regressions with variance clustered on cohorts, studies or shared control groups applied a small-sample correction and used t-based confidence intervals and tests with G−1 degrees of freedom, where G is the number of clusters. All tests were two-sided. Interaction estimates are reported as HRs per +1 s.d. of weight, except for the ordinal matched-quartile trend, which is per heavier quartile. Weighted or conventional Kaplan–Meier medians were used for direct lifespan contrasts as specified. Median-lifespan contrasts summarize absolute benefits across weight strata; individual-level effect modification was assessed using the corresponding Cox interaction or trend. Primary inference rests on the unselected all-experiment interaction; successful-experiment, matched-quartile, comparator, experiment-level weight-change, developmental, cross-study and external-population analyses are complementary. Secondary P values were not adjusted for multiple comparisons. Analyses were performed in R 4.5.1 and Python 3.13 using survival 3.8-3, pandas 2.3.2, NumPy 2.3.2, SciPy 1.18.0, lifelines 0.30.3 and Matplotlib 3.10.5.

## Data availability

Individual ITP body weights and survival are available from the Mouse Phenome Database (projects ITP1 and ITP2004–ITP2021) and GeneNetwork. Data for the external cohorts are available from the original studies: Diversity Outbred cohort^17^ and olfactory-exposure cohort^18^. Supplementary Data 1–10 provide the harmonized mouse-level data and supporting tables. The cross-study corpus is from DrugAge and subsequent ITP reports. Founder-strain data are available from the Mouse Phenome Database Reed1 project (measures 32603 and 32680) and measure 10331.

## Code availability

Code and input data to reproduce the analyses and figures are available at https://github.com/QuantCell/body-weight-geroprotector-efficacy.

## Acknowledgements

The authors acknowledge the NIA Interventions Testing Program investigators for coordinated multi-site data collection and public release, and the investigators who generated the GeneNetwork and DRiDO resources.

## Author contributions

PB conceived the study and performed the analyses, PB and LP interpreted the results and wrote the manuscript.

## Ethics approval

This secondary analysis used publicly available animal-study data; no new animal experiments were performed.

## Extended Data Tables

**Extended Data Table 1.**
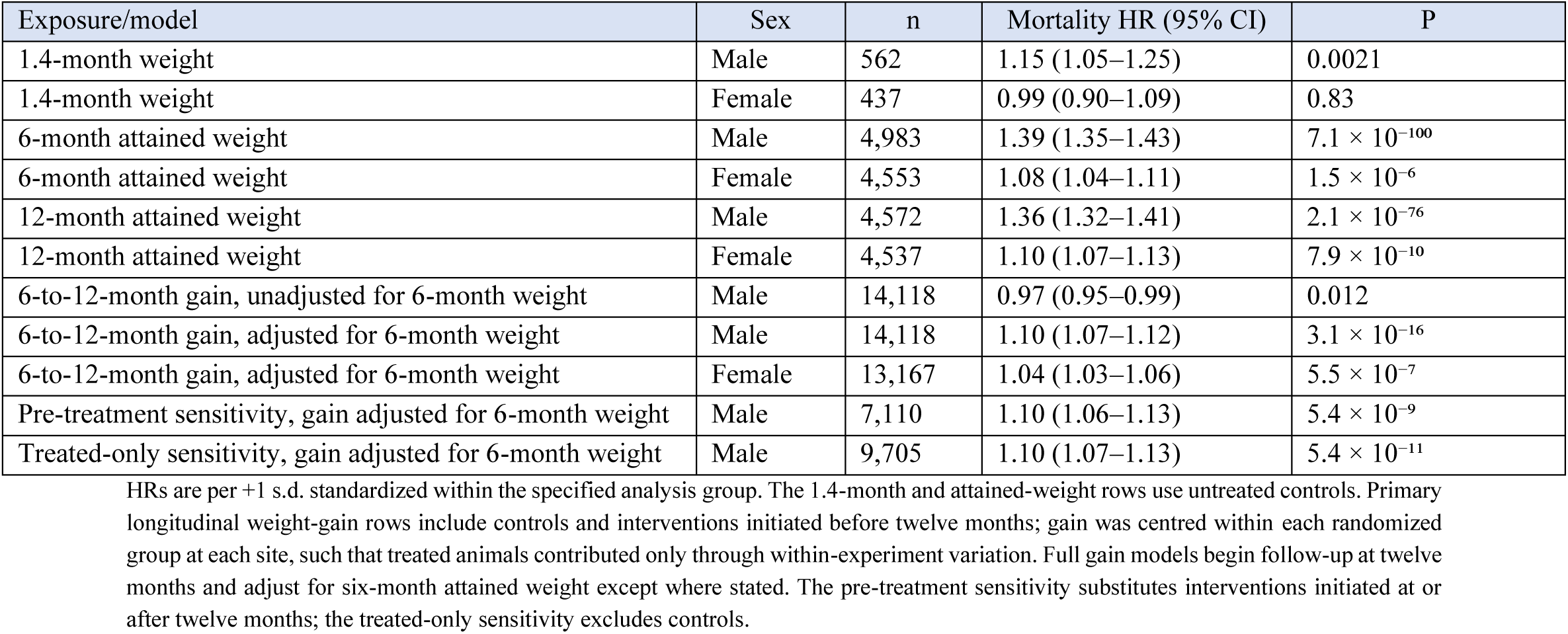
Developmental and age-specific weight–mortality estimates.

| Exposure/model | Sex | n | Mortality HR (95% CI) | P |
|---|---|---|---|---|
| 1.4-month weight | Male | 562 | 1.15 (1.05–1.25) | 0.0021 |
| 1.4-month weight | Female | 437 | 0.99 (0.90–1.09) | 0.83 |
| 6-month attained weight | Male | 4,983 | 1.39 (1.35–1.43) | 7.1 × 10<sup>−100</sup> |
| 6-month attained weight | Female | 4,553 | 1.08 (1.04–1.11) | 1.5 × 10<sup>−6</sup> |
| 12-month attained weight | Male | 4,572 | 1.36 (1.32–1.41) | 2.1 × 10<sup>−76</sup> |
| 12-month attained weight | Female | 4,537 | 1.10 (1.07–1.13) | 7.9 × 10<sup>−10</sup> |
| 6-to-12-month gain, unadjusted for 6-month weight | Male | 14,118 | 0.97 (0.95–0.99) | 0.012 |
| 6-to-12-month gain, adjusted for 6-month weight | Male | 14,118 | 1.10 (1.07–1.12) | 3.1 × 10<sup>−16</sup> |
| 6-to-12-month gain, adjusted for 6-month weight | Female | 13,167 | 1.04 (1.03–1.06) | 5.5 × 10<sup>−7</sup> |
| Pre-treatment sensitivity, gain adjusted for 6-month weight | Male | 7,110 | 1.10 (1.06–1.13) | 5.4 × 10<sup>−9</sup> |
| Treated-only sensitivity, gain adjusted for 6-month weight | Male | 9,705 | 1.10 (1.07–1.13) | 5.4 × 10<sup>−11</sup> |

HRs are per +1 s.d. standardized within the specified analysis group. The 1.4-month and attained-weight rows use untreated controls. Primary longitudinal weight-gain rows include controls and interventions initiated before twelve months; gain was centred within each randomized group at each site, such that treated animals contributed only through within-experiment variation. Full gain models begin follow-up at twelve months and adjust for six-month attained weight except where stated. The pre-treatment sensitivity substitutes interventions initiated at or after twelve months; the treated-only sensitivity excludes controls.

**Extended Data Table 2.** Lifespan increase by within-cohort quartile of treatment-associated weight change.

| Sex | Quartile | Experiments | Median weight change | Mean lifespan change | Median lifespan change | Experiments with longer lifespan |
|---|---|---|---|---|---|---|
| Male | Q1 greatest loss | 24 | −4.8% | +10.9% | +11.6% | 83.3% |
| Male | Q2 | 16 | −0.5% | +4.1% | +3.3% | 75.0% |
| Male | Q3 | 15 | +0.4% | +4.4% | +4.8% | 80.0% |
| Male | Q4 least loss or gain | 19 | +1.5% | +2.7% | +3.7% | 78.9% |
| Female | Q1 greatest loss | 22 | −11.3% | +4.6% | +2.1% | 81.8% |
| Female | Q2 | 15 | −4.0% | +1.4% | −1.0% | 46.7% |
| Female | Q3 | 14 | −0.5% | +1.5% | −0.7% | 42.9% |
| Female | Q4 least loss or gain | 17 | +2.1% | +0.6% | −0.4% | 35.3% |

Quartiles were assigned within cohort and sex among cohorts with at least four eligible experiments. Of 163 contrasts in the continuous analysis, 142 contributed here (74 male contrasts in 13 cohorts and 68 female contrasts in 12 cohorts). All directions of lifespan change were retained. Q1 denotes the greatest relative weight reduction within each cohort, not a common absolute cut-off. Cohort-clustered trends per quartile toward greater weight loss were +2.56 percentage points in males (95% CI 0.53–4.60; P = 0.018) and +1.26 in females (−0.28 to 2.80; P = 0.098).

**Extended Data Table 3.**
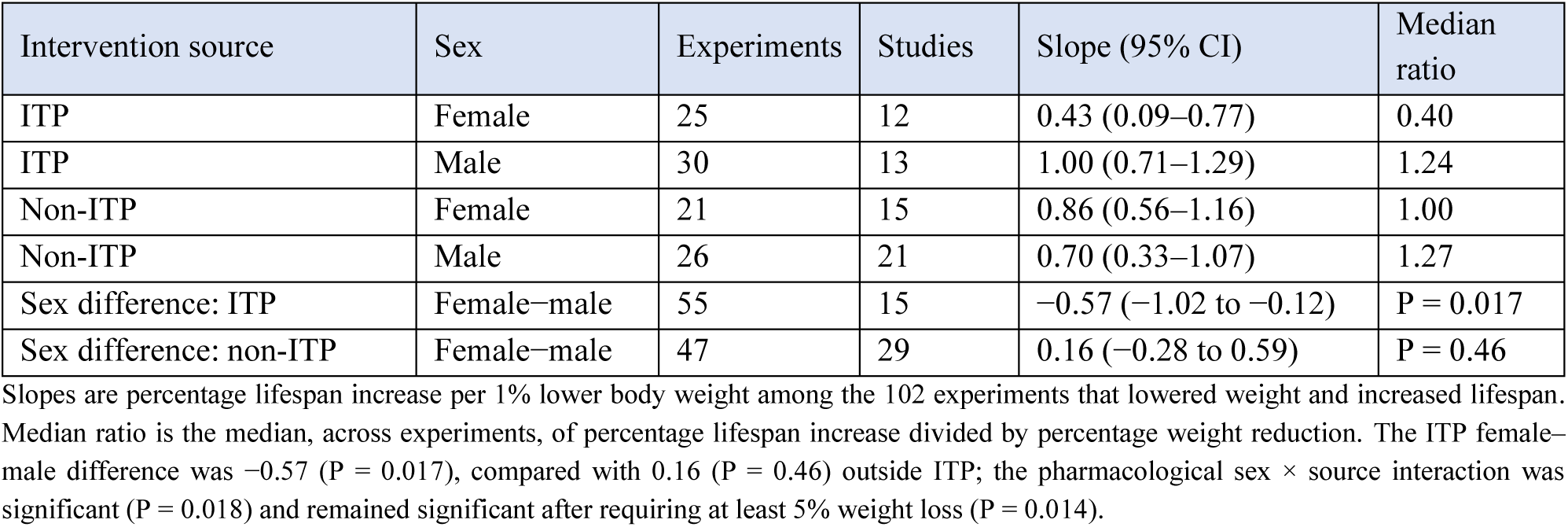
Weight reduction and lifespan increase by study source and sex.

| Intervention source | Sex | Experiments | Studies | Slope (95% CI) | Median ratio |
|---|---|---|---|---|---|
| ITP | Female | 25 | 12 | 0.43 (0.09–0.77) | 0.40 |
| ITP | Male | 30 | 13 | 1.00 (0.71–1.29) | 1.24 |
| Non-ITP | Female | 21 | 15 | 0.86 (0.56–1.16) | 1.00 |
| Non-ITP | Male | 26 | 21 | 0.70 (0.33–1.07) | 1.27 |
| Sex difference: ITP | Female−male | 55 | 15 | −0.57 (−1.02 to −0.12) | P = 0.017 |
| Sex difference: non-ITP | Female−male | 47 | 29 | 0.16 (−0.28 to 0.59) | P = 0.46 |

Slopes are percentage lifespan increase per 1% lower body weight among the 102 experiments that lowered weight and increased lifespan. Median ratio is the median, across experiments, of percentage lifespan increase divided by percentage weight reduction. The ITP female–male difference was −0.57 (P = 0.017), compared with 0.16 (P = 0.46) outside ITP; the pharmacological sex × source interaction was significant (P = 0.018) and remained significant after requiring at least 5% weight loss (P = 0.014).

**Extended Data Table 4.** Median lifespan by pre-intervention weight quartile in female Diversity Outbred mice.

| Pre-intervention weight quartile | Mean weight | AL median lifespan | Pooled CR median lifespan | Increase versus AL |
|---|---|---|---|---|
| Lowest-weight | 24.8 g | 877 days (n = 47) | 973 days (n = 93) | +96 days (+10.9%) |
| Second | 29.8 g | 825 days (n = 47) | 974 days (n = 93) | +149 days (+18.1%) |
| Third | 34.6 g | 779 days (n = 47) | 931 days (n = 92) | +152 days (+19.5%) |
| Highest-weight | 42.1 g | 714 days (n = 47) | 946 days (n = 93) | +232 days (+32.5%) |

The analysis included 559 Diversity Outbred females with body weight measured within 45 days of the 180-day landmark and surviving beyond it. Weight quartiles were formed separately within the *ad libitum* and pooled caloric-restriction groups; the caloric-restriction groups were pooled for display. Medians are Kaplan–Meier lifespan from birth, and increase is the pooled-restriction minus *ad libitum* median within each quartile. The pooled treatment-by-quartile trend was HR 0.90 (95% CI 0.76–1.05; P = 0.18). Weight predicted mortality among *ad libitum* controls (HR 1.21 per s.d., 1.04–1.41; P = 0.015) but not under restriction (HR 1.05, 0.95–1.17). AL, *ad libitum*; CR, caloric restriction.

**Extended Data Table 5.** Weight–mortality gradients in ITP controls and external individual-level cohorts.

| Cohort and weight measure | Sex | Mice | Dead by day 500 | Mortality HR per s.d. (95% CI) | P |
|---|---|---|---|---|---|
| ITP controls, six-month weight | Male | 4,983 | 16.0% | 1.39 (1.35–1.43) | 7.1 × 10<sup>−100</sup> |
| ITP controls, six-month weight | Female | 4,553 | 2.4% | 1.08 (1.04–1.11) | 1.5 × 10<sup>−6</sup> |
| Diversity Outbred *ad libitum* controls, pre-intervention weight | Female | 188 | 12.2%* | 1.21 (1.04–1.41) | 0.015 |
| Olfactory-exposure cohort, no-odour controls, six-month weight | Female | 59 | 32.1% | 1.52 (1.14–2.02) | 0.0042 |
| Olfactory-exposure cohort, no-odour controls, six-month weight | Male | 60 | 13.3% | 1.18 (0.91–1.52) | 0.21 |

HRs are per +1 s.d. of weight standardized within the analysis group (ITP: cohort × site; Diversity Outbred: *ad libitum* group; olfactory-exposure cohort: sex × exposure × genotype), with follow-up from the weighing landmark. ITP models were stratified by cohort and site, and olfactory-exposure models by genotype. Day-500 mortality is the Kaplan–Meier estimate in all controls, censoring removals; \*the Diversity Outbred source does not record removal reasons, so every animal counts as an event (ITP females on that basis: 3.0%). Among all 170 olfactory-exposure females, six-week weight was not associated with mortality after adjustment for six-month weight (HR 1.01, 0.82–1.24; model stratified by exposure and genotype). Weight–mortality HR intervals are model-based for all cohorts.

**Extended Data Table 6.** Sex-specific survival context in ITP and matched external controls.

| Population / source | Metric | Male | Female | Female–male difference |
|---|---|---|---|---|
| ITP pooled controls (all 17 cohorts) | Kaplan–Meier median survival | 800 days (n = 5,352) | 886 days (n = 4,833) | +86 days |
| Independent UM-HET3;<sup>22</sup> 5K54 | Reported mean survival | 831 days<br>(n = 14) | 836 days<br>(n = 16) | +5 days |
| Independent 5LG6 cohort;<sup>23</sup><br>C57BL/6J | Reported median survival | 804 days<br>(n = 40) | 865 days<br>(n = 45) | +61 days |
| ITP pooled controls (all 17 cohorts) | Kaplan–Meier mortality by day 500 | 16.0%<br>(n = 5,352) | 2.4%<br>(n = 4,833) | −13.6 percentage points |
| Independent Diversity Outbred *ad libitum*<sup>17</sup> | Kaplan–Meier mortality by day 500 | – | 12.2% (n = 188) | – |

ITP medians and day-500 mortality were calculated from individual untreated controls. External rows were selected to match UM-HET3 ancestry and 5LG6 chow separately.

**Extended Data Table 7.** ITP control survival relative to published non-ITP control groups.

| Specification | Sex | ITP groups | ITP median (d) | Non-ITP groups | Non-ITP median (d) | Non-ITP reaching ITP median |
|---|---|---|---|---|---|---|
| Unique control groups | Female | 18 | 891 | 17 | 815 | 0 of 17 |
| Unique control groups | Male | 18 | 801 | 20 | 814 | 10 of 20 |
| Study-level median | Female | 13 | 893 | 15 | 724 | 0 of 15 |
| Study-level median | Male | 14 | 796 | 18 | 794 | 9 of 18 |
| Median endpoints only | Female | 18 | 891 | 9 | 847 | 0 of 9 |
| Median endpoints only | Male | 18 | 801 | 9 | 912 | 5 of 9 |
| Published 2000 or later | Female | 18 | 891 | 12 | 842 | 0 of 12 |
| Published 2000 or later | Male | 18 | 801 | 14 | 835 | 8 of 14 |

Control-group survival after collapsing intervention experiments that share a study and reported control lifespan, because ITP publications commonly report several experiments against one control; the underlying 221 experiments are listed in Supplementary Data 10. ITP status required membership of an Interventions Testing Program cohort; UM-HET3 studies conducted outside the programme were classified as non-ITP. For the unique-group, study-level, median-endpoint and post-2000 specifications, two-sided Mann–Whitney P values were 2.5 × 10<sup>−6</sup>, 1.5 × 10<sup>−5</sup>, 2.9 × 10<sup>−4</sup> and 3.6 × 10<sup>−5</sup> for females and 0.51, 0.86, 0.29 and 0.15 for males. Values in this table are medians across published control-group lifespan estimates, which follow the endpoint reported by each source study except in the median-endpoint sensitivity.

**Extended Data Table 8.** Male body weight in the four UM-HET3 founder strains.

| Founder strain | Male weight |
|---|---|
| BALB/cByJ | 32.0 g |
| C57BL/6J | 30.8 g |
| C3H/HeJ | 26.7 g |
| DBA/2J | 28.1 g |
| Four-founder mean | 29.4 g |

Unadjusted male means from Mouse Phenome Database Reed1 measure 32603; corresponding mean ages were approximately 30.3–31.5 weeks from measure 32680 (n = 10–12 per strain).<sup>11</sup> Mice were individually housed with *ad libitum* access to Teklad Rodent Diet 8604. The four-founder mean is the unweighted mean of the strain estimates.

